# Effects of Historical Coinfection on Host Shift Abilities of Exploitative and Competitive Viruses

**DOI:** 10.1101/105114

**Authors:** Sonia Singhal, Paul E. Turner

**Affiliations:** Department of Ecology and Evolutionary Biology, Yale University, P. O. Box 208106, New Haven, Connecticut 06520, USA; Graduate Program in Microbiology, Yale School of Medicine, New Haven, Connecticut 06520, USA

**Keywords:** RNA virus, ecological history, host shift, multiplicity of infection

## Abstract

Rapid evolution contributes to frequent emergence of RNA viral pathogens on novel hosts. However, accurately predicting which viral genotypes will emerge has been elusive. Prior work with lytic RNA bacteriophage f6 (family Cystoviridae) suggested that evolution under low multiplicity of infection (MOI; proportion of viruses to susceptible cells) selected for greater host exploitation, while evolution under high MOI selected for better intracellular competition against co-infecting viruses. We predicted that phage genotypes that experienced 300 generations of low MOI ecological history would be relatively advantaged in growth on two novel hosts. We inferred viral growth through changes in host population density, specifically by analyzing five attributes of growth curves of infected bacteria. Despite equivalent growth of evolved viruses on the original host, low MOI evolved clones were generally advantaged relative to high MOI clones in exploiting novel hosts. We also observed genotype-specific differences in clone infectivity: High fitness genotypes on the original host also performed better on novel hosts. Our results indicated that traits allowing greater exploitation of the original host correlated positively with performance on novel hosts. Based on infectivity differences of viruses from high versus low MOI histories, we suggest that prior MOI selection can later affect emergence potential.

## Introduction

RNA viruses are able to evolve rapidly due to high mutation rates, large population sizes, and short generation times (Wasik and Turner 2013). These characteristics seem to foster successful emergence of RNA viruses as pathogens on novel host types: In humans, RNA viruses are overrepresented among emerging and re-emerging pathogens (Woolhouse and Gowtage-Sequeria 2005). Examples of recent emerging diseases caused by RNA viruses include AIDS (Barré-Sinoussi *et al*. 1983, Levy *et al*. 1984), influenza (e.g., Ginsberg *et al*. 2009, Jordan *et al*. 2009), Ebola hemorrhagic fever (Baize *et al*. 2014, Dixon and Schafer 2014), and Zika fever (e.g., Duffy *et al*. 2009, Faye *et al*. 2014).

However, not all genotypes in a viral population should be equally capable of growing robustly in a novel host. Consistent with this idea, individual viral genotypes vary in their fitness across host types (Turner *et al*. 2010, Deardorff *et al*. 2011, Lalić *et al*. 2011, Vale *et al*. 2012, Ford *et al*. 2014, Burmeister *et al*. 2016), indicating that certain genotypes are better poised to infect novel hosts. These genotypes are shaped both by spontaneous mutation and selective pressures previously experienced during evolution. In some cases, prior selective pressures may have pleiotropic or correlated effects that permit infection of a novel host. For example, adaptation to alternating cell types permitted the evolution of “generalist” vesicular stomatitis virus (VSV) strains that were better able to infect cell types they had not previously experienced (Turner *et al*. 2010). Even adaptation on a single host can fortuitously permit exploitation of other, distantly related hosts. For example, evolution of VSV on a single cell type with an intact innate immune response seemed to foster the evolved virus’s ability to infect other hosts with similar immune responses (Wasik *et al*. 2016). Prior ecological history can thus affect viral performance in new host environments.

In addition to external pressures due to the host environment, inherent population dynamics can impose selective pressures. A classic example of how selection can change as a result of population dynamics is frequency-dependent selection, in which the fitness of a genotype changes depending on how rare or common it is relative to other genotypes (e.g., Lewontin 1958, Clarke and O’Donald 1964, Smouse 1976). Because viruses rely on susceptible host cells for replication, an important aspect of viral population dynamics is the multiplicity of infection (MOI), the ratio of viral particles to susceptible cells (Gutiérrez *et al*. 2010, Lancaster and Pfeiffer 2012). MOI can change across viral generations. For example, infectious viral diseases seem to be initiated by only few viral particles (Gutiérrez *et al*. 2012), indicating low initial MOI. Over time, if viral replication locally outpaces reproduction of susceptible host cells, MOI will correspondingly increase. Depletion of the local cell population or transmission to a new multicellular host then induces bottlenecks, often severe ones, and a concomitant decrease in MOI (Gutiérrez *et al*. 2012). Subsequent expansion of the virus population may again increase MOI.

Changes in MOI can alter the competitive environment for a viral genotype. At MOI >1 (i.e., average of more than one viral particle per susceptible host cell), it becomes likely that two or more viruses will co-infect the same cell. Co-infecting viruses may compete within the cell for resources such as amino acids and access to replicative machinery. Intracellular competition can lead to the evolution of “cheating” strategies among viruses. For example, viruses that specialize in sequestering and assembling proteins may take advantage of the proteins encoded by co-infecting genotypes that instead maximize transcription (Turner and Duffy 2008). Cheater viruses may even evolve reduced genome sizes by deleting essential genes, enabling faster replication despite their reliance on fully functional genotypes (Huang and Baltimore 1970, Huang 1973, Williams *et al*. 2016).

Despite the importance of MOI in viral population dynamics, its role in the subsequent performance of viral lineages has rarely been explored. A previous study evolved f6 Cystovirus, a lytic bacteriophage with a segmented RNA genome, under high MOI (average of 5 viral particles/cell; hereafter H lineages) or low MOI (average of 0.002 viral particles/ cell; hereafter L lineages) (Turner and Chao 1998). MOI was controlled at each transfer by mixing viruses with *Pseudomonas syringae* pathovar *phaseolicola* host bacteria at the treatment MOI, allowing them to adsorb (attach to the host), and then plating them. Plaques were harvested for the next round of adsorption on naïve (non-coevolving) host bacteria. Fitness evaluations after 250 generations of evolution indicated that the dominant selective pressure differed depending on the MOI treatment. Clones from L lineages had increased their ability to exploit the bacterial host with respect to the ancestral phage. Clones from H lineages, however, demonstrated MOI-dependent fitness: Performance relative to the ancestor increased at high MOI but was reduced at low MOI, suggesting that selection for intracellular competition can interfere with generalized adaptations to exploit host cells (Turner and Chao 1998; see also Turner and Chao 1999, 2003). By generation 300, however, fitness differences between low MOI and high MOI evolved clones were no longer detectable, even though the viruses from low MOI and high MOI treatments had diverged from one another genetically (McBride *et al*. 2008, Dennehy *et al*. 2013). The low MOI and high MOI evolved clones thus represent genetically different viruses, molded by selection in each environment, that can be harnessed to test how MOI ecological history affects viral performance on novel hosts.

### Experimental overview

In the laboratory, growth of ɸ6 is supported on the bacterium *P. phaseolicola*. We challenged ɸ6 clones, evolved for 300 generations under low or high MOI, to grow on two novel host bacteria, *P. syringae* pathovars *tagetis* and *savastanoi*. These hosts are permissive for growth of wild type ɸ6 (Duffy *et al*. 2006); we define them as “novel” because the evolved clones had not encountered them in their recent selective history. We used an automated, high-throughput method of inferring viral fitness by measuring changes in the growth of infected bacteria over time, compared to the typical logarithmic growth of uninfected bacteria (Fig. 1; see Methods and Turner *et al*. 2012). In particular, these assays yielded estimates of five variables indicating the rate and extent of a virus’s ability to kill the bacterial host population (Fig. 1, Table 1). Lower values of these variables indicated better viral performance on the host.

**Figure 1:**
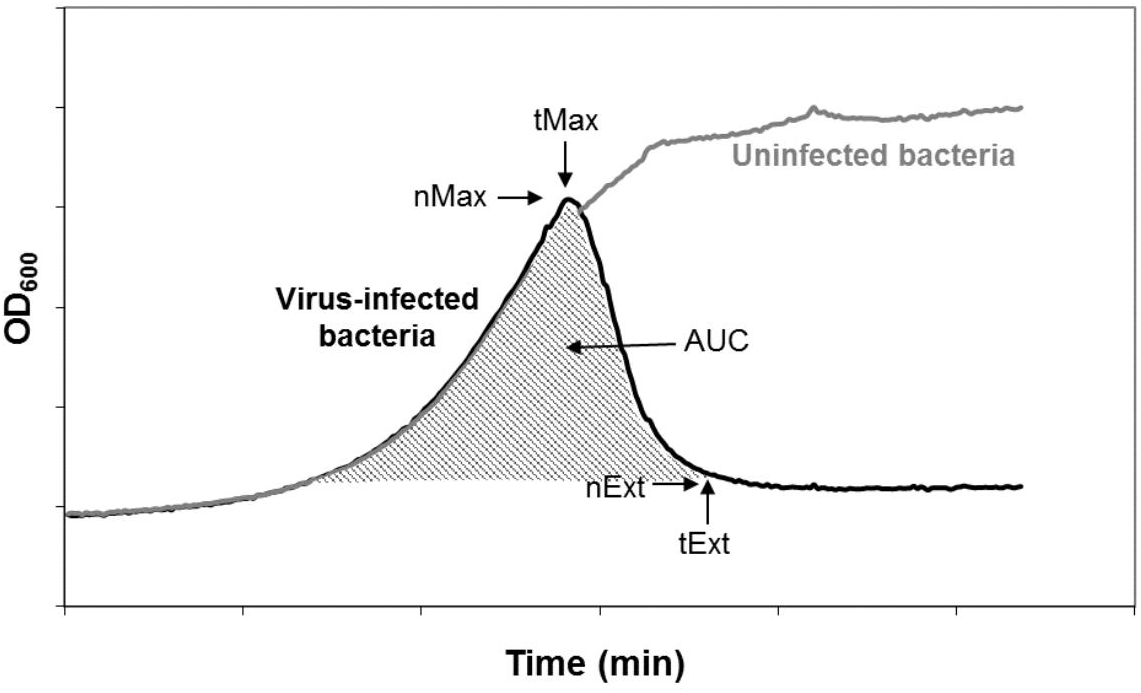
**Example growth curves of uninfected and phage-infected bacterial cultures**. Bacterial growth was estimated by measuring changes in optical density (OD_600_) over time. Viral growth performance was inferred from deviations from the logarithmic growth pattern of uninfected bacteria using 5 measures: maximum bacterial density (nMax), time of maximum density (tMax), bacterial density at extinction (nExt), time of bacterial extinction (tExt), and area under the growth curve (AUC).

**Table 1:**
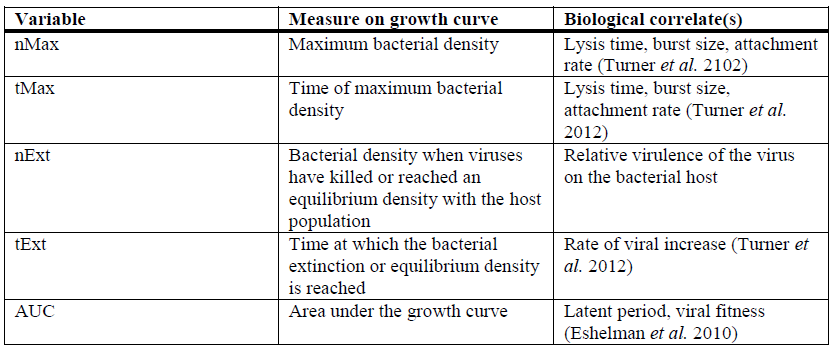
**Measures extracted from growth curves of phage-infected bacteria**. Lower values of nMax, tMax, and tExt indicate that a virus is able to infect and kill host cells more rapidly (via shorter lysis time, faster attachment rate, or larger burst size, Turner *et al*. 2012). AUC gives a holistic measure of viral fitness, with a lower AUC (less cumulative bacterial growth) corresponding to higher viral fitness (our measure is the inverse of the virulence index described in Eshelman *et al*. 2010). Viruses that are relatively better at driving their host to extinction will have a low nExt (relatively betters growth on that host), while viruses that settle into an equilibrium density with their host will have a higher nExt (relatively poorer growth on that host). See Figure 1 for details of how variables relate to dynamics of bacterial growth.

| Variable | Measure on growth curve | Biological correlate(s) |
| --- | --- | --- |
| nMax | Maximum bacterial density | Lysis time, burst size, attachment rate (Turner *et al.* 2102) |
| tMax | Time of maximum bacterial density | Lysis time, burst size, attachment rate (Turner *et al.* 2012) |
| nExt | Bacterial density when viruses have killed or reached an equilibrium density with the host population | Relative virulence of the virus on the bacterial host |
| tExt | Time at which the bacterial extinction or equilibrium density is reached | Rate of viral increase (Turner *et al.* 2012) |
| AUC | Area under the growth curve | Latent period, viral fitness (Eshelman *et al.* 2010) |

## Methods

### Strains and culture conditions

All viruses in this study were derived from wild type phage ɸ6 (American Type Culture Collection [ATCC] #21781-B1). The specific viral clones originated from a prior experiment (Turner and Chao 1998) in which three lineages were each evolved at low MOI (L lineages) or high MOI (H lineages). Montville *et al*. (2005) isolated ten clones from each lineage after 300 generations of evolution (total of 30 clones from the L lineages and 30 clones from the H lineages). The current study examined eleven of these clones as test phages (designated L1.7, L2.4, L2.6, L3.2, L3.8; H1.1, H1.3, H2.4, H2.10, H3.5, and H3.9; see also Dennehy *et al*. 2013).

We examined growth of the viral clones on the typical laboratory host for ɸ6, *P. phaseolicola* strain HB10Y (ATCC #21781) and two challenge hosts, *P. tagetis* and *P. savastanoi* (kindly provided by G. Martin, Cornell University, Ithaca, New York; Martin strain numbers 2230 and 2237, respectively). Bacterial cultures were initialized from frozen stock by streaking onto 1.5% agar plates made from LC medium (Luria-Bertani broth at pH 7.5) and incubating for 48 hours at 25°C to obtain visible colonies. An individual colony was then randomly chosen, placed in liquid LC medium, and incubated with shaking at 25°C for 2 hours to obtain a stationary-phase bacterial culture.

Viral lysates were prepared from frozen stocks of virus by mixing a diluted sample of the virus stock with 200 overlaid on a 1.5% LC agar base and incubated overnight at 25°C to obtain visible plaques. Plaques were collected in 3 mL of liquid LC medium and filtered (0.22 fL pore, Durapore, Millipore) to remove bacterial cells. Lysates were stored in 4:6 (v/v) glycerol: LC at −20°C.

### Measuring bacterial growth curves

Growth curves of phage-infected and uninfected bacteria were obtained by culturing bacteria in 200 fL of LC medium in transparent, flat-bottomed 96-well plates. The plates were placed in a microplate reader (model Infinite F500, Tecan) that measured the optical density at 600 nm (OD600) every 5 minutes for 24 hours. Between OD600 measurements, plates were incubated at 25°C with orbital shaking.

Each well of the 96-well plate contained 4×10^7^cells of stationary-phase bacteria with (infected) or without (uninfected) 400 plaque forming units (pfu) of a test phage. Each plate housed seven replicate cultures of uninfected bacteria (controls) and seven replicates of bacteria infected with each of the eleven test phages (84 culture wells total). Separate plates were used for each of the three host bacteria (three 96-well plates total). A random number generator was used to assign the locations of each replicate within the 96-well plate, and these locations were held constant across plates examining different hosts. Growth curves were repeated on two separate days to obtain 14 replicate measurements of each bacterial host infected with each of the eleven test phages.

### Extracting measures of viral growth from bacterial growth curves

Similar to Turner *et al*. (2012), five variables were estimated from the bacterial growth curves (OD_600_ data; Fig. 1, Table 1): (i) maximum bacterial density (nMax); (ii) time at which bacteria reached maximum density (tMax); (iii) time at which bacteria went extinct or reached an equilibrium density with the test phage (tExt); (iv) OD_600_ reading at time tExt (nExt); and (v) area under the growth curve (AUC).

A script (see Data repository) written in R (version 3.1.2) smoothed the growth curves by filtering the OD_600_ values at each time point through a repeated medians filter (robfilter package). Variables nMax and tMax were set as the maximum OD_600_ value of the smoothed curves and the corresponding time. Variables nExt and tExt were determined by the point where the difference in sequential OD_600_ values dropped below a threshold (set as the 50% quantile of all differences between sequential OD_600_ measurements). AUC was calculated using Riemann trapezoidal sums at 5-minute intervals. For ease of comparison across hosts with different growth rates and densities, all variables were normalized by the median value of the corresponding uninfected bacterial controls.

Further details on data processing methods are available in the Data repository. Examples of growth curves before and after processing are shown in Fig. S1.

### Statistical analyses

Data did not meet normality assumptions of parametric tests, even after log transformation of the estimated variables. For this reason, we ran non-parametric tests for conservative estimates of how different viruses affected the growth of each bacterial host. Kruskal-Wallis rank sum tests were used to assess significant differences in variables for hosts that were uninfected, infected by low MOI (L) viruses, or infected by high MOI (H) viruses. Post-hoc permutation tests (10,000 permutations) were then used to determine specifically which viral clones differed from one another. Significance levels were adjusted using a Bonferroni correction. All analyses were run in R (version 3.1.2). Scripts are available in the Data repository.

### Results

For eleven phage clones previously evolved for 300 generations at low (L) or high MOI on *P. phaseolicola*, we estimated five variables of their performance by comparing growth curves of infected bacteria to those of uninfected controls (see Methods). The medians and 95% quantile ranges of these extracted variables on each host, normalized against the median values for the uninfected controls, are shown in Fig.2 Lines connect variable estimates for better visualization of the overall patterns of similarity and difference between groups of viral clones and across host environments.

**Figure 2:**
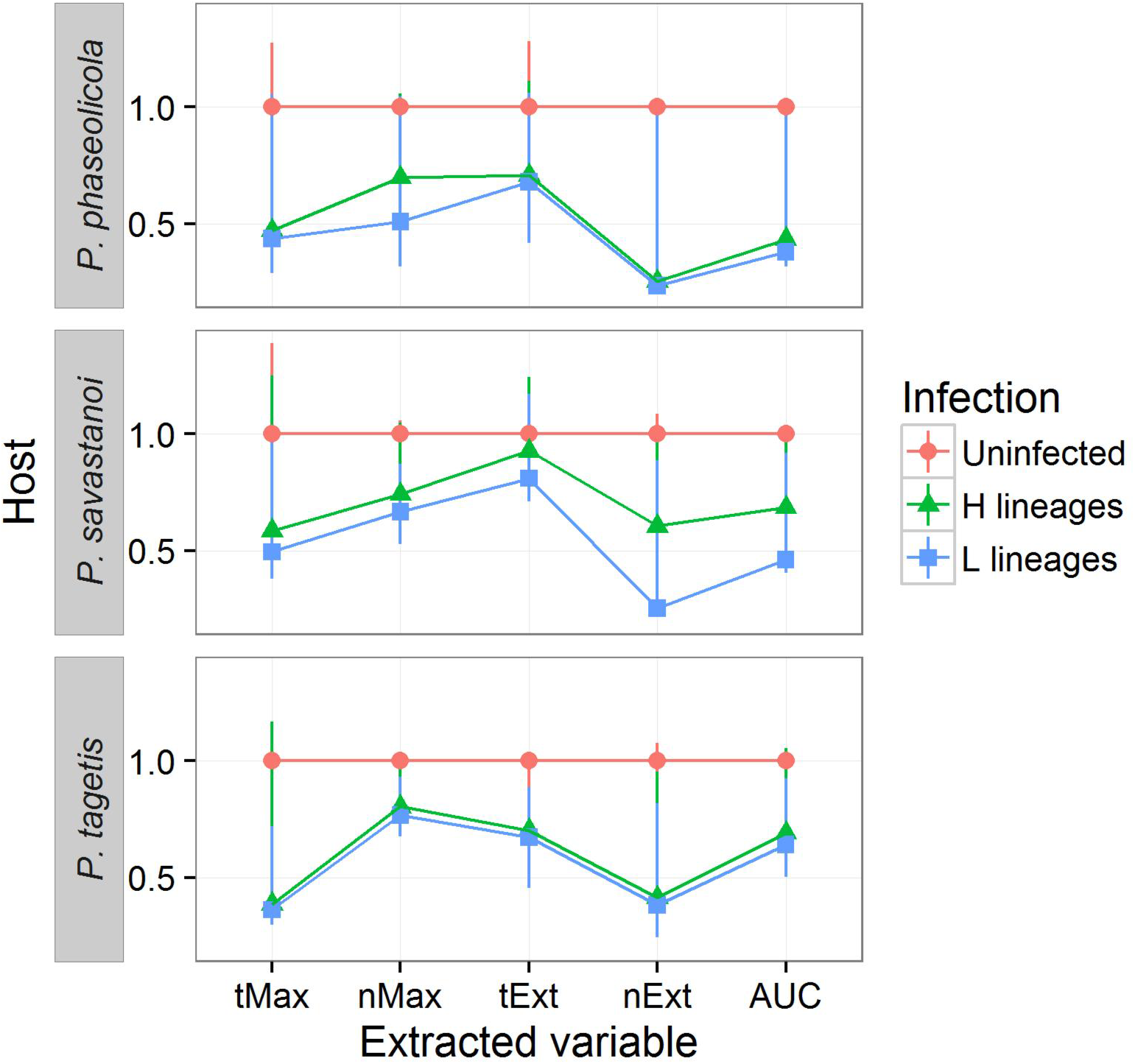
**Medians of variables extracted from growth curves of uninfected bacteria and bacteria infected with phages evolved at low MOI (L) or high MOI (H)**. All variables were normalized against the median values for uninfected bacteria. Error bars represent the 95% quantile range of the data. Lines connect variables for easier visualization of comparisons in performance between L and H viruses on each host.

To examine whether L and H clones differed in their effects on bacterial growth, we ran separate Kruskal-Wallis rank sum tests on each variable and each host. For all variables, the rank orders differed significantly based on whether the host was uninfected, infected with a clone from an L lineage, or infected with a clone from an H lineage (p << 0.001; Table S1). Post-hoc permutation tests revealed that, across hosts and variables, rankings for L clones were significantly higher than rankings for H clones (i.e., closer to a ranking of 1, or the minimum value in the dataset; Table S2). In other words, compared to the H clones, L clones tended to show lower mean values for each variable, indicating superior performance. The only variables that presented no significant differences between L and H clones were tExt on *P. tagetis*, and tExt and tMax on *P. phaseolicola*.

Clones within groups of L and H viruses varied in their performance, especially on the novel hosts (Fig. 3). However, overall rankings of viral clones were similar across hosts (Spearman’s *ρ* > 0.60, p < 0.02; with the exception of tExt on *P. tagetis*, for which correlations were not significant; Table S3). High-ranked clones on *P. phaseolicola* also tended to rank relatively well on novel hosts, while low-ranked clones from *P. phaseolicola* also performed relatively poorly on the novel hosts. This trend suggested that fitness on the novel hosts might simply be a function of fitness on the laboratory host. To examine this possibility more closely, we compared two general linear models for each variable: one model that predicted a clone’s rank solely from its rank on *P. phaseolicola*, and another model that additionally incorporated whether the virus had been evolved under low or high MOI (Table S4). For two variables on *P. tagetis* (nMax and tMax) and all variables on *P. savastanoi*, the model that included MOI treatment explained more of the variation in rankings than did the model that only considered ranking on the original host (p ≤ 0.02 in all cases mentioned here).

**Figure 3:**
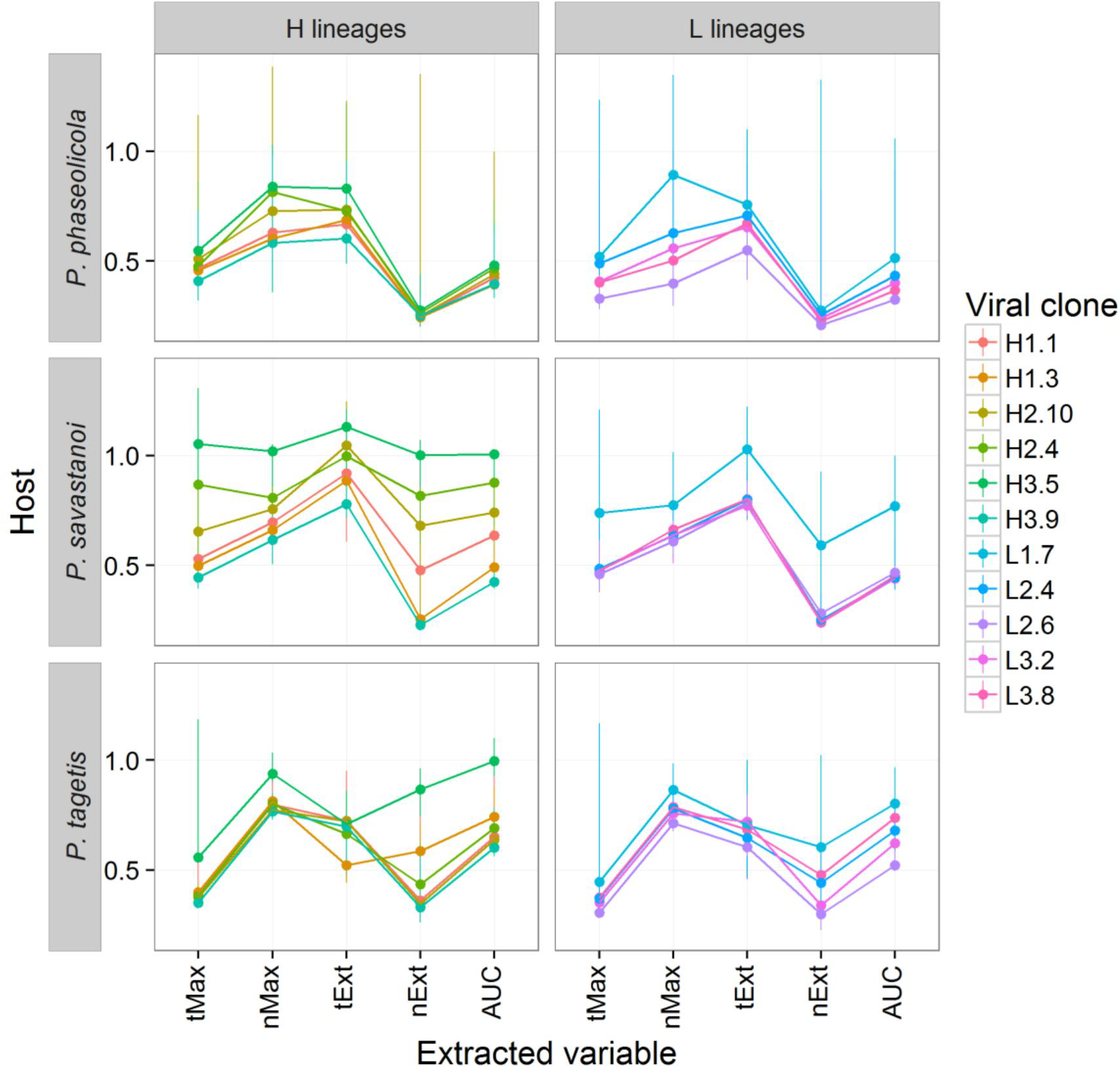
**Medians of variables extracted for each evolved phage clone on three hosts**. All variables were normalized against the mean values for uninfected bacteria. Error bars represent the 95% quantile range of the data. Lines connect variables for easier visualization of comparisons in performance between L and H viruses on each host.

On a single host, rankings of the individual viral clones were highly correlated across extracted variables (Table S5): Clones that performed relatively well (closer to a ranking of 1) in one variable tended to perform well in the others (Spearman’s *ρ* between 0.88 – 0.99, p << 0.001; except for tExt on *P. tagetis*, for which correlations were not significant). Host effects on the growth of the viral clones are apparent in the similarities in relationships among variables from different clones on the same host (Fig. 2, Fig. 3). Corroborating this observation, Spearman correlation coefficients of each variable exhibited a narrower range and closer correlations within a single host (excluding tExt on *P. tagetis, ρ*> 0.88, p << 0.001; Table S5) than across hosts (0.56 < *ρ*< 0.86, excluding tExt on *P. tagetis*, p ≤ 0.027; Table S3).

### Discussion

Multiplicity of infection (MOI) affects viral population dynamics, and relatively low versus high MOI can exert differing selective pressures on viral populations (Huang and Baltimore 1970, Huang 1973, Horiuchi 1973, Turner and Chao 1998, 1999, 2003, Gutiérrez *et al*. 2010, Lancaster and Pfeiffer 2012, Williams *et al*. 2016). However, the effects of historical MOI selection on a virus’s subsequent performance on alternative hosts are not widely studied. A prior study evolved the phage ɸ6 on the original *P. phaseolicola* host under low or high MOI. Here, we examined how 300 generations of MOI selection affected the ability of viral clones to infect two novel hosts: *P. tagetis* and *P. savastanoi*. We used a high-throughput approach that tracked changes in the growth of infected bacterial cultures over time (see also Turner *et al*. 2012). From these data, we estimated five variables to describe the effects of viral infection on host growth, compared to uninfected bacterial controls. These variables indicated how quickly (tMax, tExt) and how completely (nMax, nExt, AUC) a viral clone reduced host population density. Viruses with lower tMax and tExt values killed hosts more quickly (bacterial density was reduced earlier); similarly, lower values of nMax, nExt, and AUC indicated that a virus was better at reducing peak, final, and overall bacterial densities, respectively. In phage ɸ6, these measures were previously shown to correlate strongly with the results of traditional fitness assays conducted on agar plates (Turner *et al*. 2012).

At high MOI, virus co-infection of host cells is more likely, which can produce frequent intracellular competition for limited host resources among co-infecting genotypes. In contrast, at low MOI virus co-infection is relatively rare, and intracellular competition should be less important as a selective pressure. Prior studies indicated that 250 generations of low versus high MOI conditions were consequential to the fitness of phage Φ6 (Turner and Chao 1998). Low MOI evolved viruses increased fitness on *P. phaseolicola* relative to the wild type ancestor in both low and high MOI environments, suggesting that selection for increased host exploitation was sufficient for generalized improvement on that host. On the other hand, high MOI evolved viruses showed increased fitness in their selective environment (high MOI) but constrained performance at low MOI, suggesting that selection for intracellular competition reduced the general ability to evolve host exploitation (Turner and Chao 1998, 1999). Importantly, after 300 generations of MOI selection, there were no fitness differences detectable by traditional assays between groups of viruses evolved at low and high MOI on *P. phaseolicola* (Figure S2; Montville *et al*. 2005, McBride *et al*. 2008, Dennehy *et al*. 2013). The current study tested whether this historical MOI selection would nonetheless influence the performance of Φ6 viruses on challenge hosts.

We confirmed that historical selection under low or high MOI could affect the subsequent performance of viral clones on novel bacterial hosts. As indicated from their significantly lower mean values of tMax on both challenge hosts, and of tExt on one challenge host, clones evolved at low MOI generally killed the novel hosts more quickly than their high MOI evolved counterparts. L clones also reduced host density more completely than H clones, indicated by their significantly lower mean values of nMax, nExt, and AUC on both novel hosts. Our results confirm that prior selection at low versus high MOI on the original host affected subsequent growth (emergence) of the viruses on two novel hosts.

Prior work shows that the evolved clones in our study have diverged genetically, but they no longer differ in fitness across MOI environments when infecting the typical *P. phaseolicola* host (Montville *et al*. 2005, McBride *et al*. 2008, Dennehy *et al*. 2013). However, in the current study, L clones were, on average, advantaged over H clones in nMax, nExt, and AUC on *P. phaseolicola.* This difference may be explained by the performance of one especially high-fitness clone, L2.6, which permutation tests revealed to be significantly different (i.e., lower values of each variable) from many of the other clones in this study (Table S6). Another possibility is that the current method of tracking changes in optical density is useful for elucidating the multiple fitness components that gauge the success of viral infection in a 24 hour period. Traditional fitness assays ignore these components for the purposes of convenience and simply resolve overall differences in viral productivity. In particular, traditional fitness assays cannot reveal how virus infection affects the density of the host bacterial population over time. It is precisely such metrics of host density (nMax, nExt, AUC) that were observed to differ between L and H clones on the typical *P. phaseolicola* host. Nevertheless, our analysis showed that the emergence success of viral clones was not trivially due to growth differences on the typical host: Generalized linear models showed that the ranked performance of viral clones on the novel hosts were best explained when prior MOI history was included.

The extracted variables could be used more specifically to identify what aspects of viral growth changed on the novel hosts. For example, the variables tMax and tExt reflect how quickly a virus infects a cell and causes it to release progeny (burst). These variables tended to be lower in value (i.e., better for viral fitness) on *P. tagetis* than on *P. savastanoi* (Table S7). These results were consistent with data from classic burst experiments (measurement of changes in viral titer over time) when wild type ɸ6 was assayed on each of the novel host types: Lysis time on *P. tagetis* was shorter than on *P. savastanoi* (Figure S3). In contrast, variables nMax and nExt indicate the proportion of host cells killed by the virus. These estimates were typically lowest on *P. phaseolicola* and highest on *P. tagetis* (Table S7). A high nExt (host extinction density), in particular, indicates that the host environment is relatively difficult (less permissive) for viral infection. Hosts with a high nExt are not killed as completely by the infecting virus, suggesting that such host/virus combinations may be more likely to achieve an equilibrium. Future work could examine co-evolution between Φ6 viruses and the host bacteria used in the current study to test whether one or more of the extracted variables usefully predicts bacteria/phage evolutionary dynamics and the possibility that a population will achieve a stable equilibrium.

Our main conclusion – that evolution under low MOI can promote viral performance on novel hosts – is corroborated in studies with poliovirus, an RNA virus that infects humans. Stern *et al*. (2014) simulate adaptation of poliovirus after transmission to a novel host type. The poliovirus particles that established the transmission event were assumed to be at an evolutionary equilibrium, following evolution at either low MOI (0.1 viral particles/cell) or high MOI (>10 viral particles/cell). Transmission then imposed a severe bottleneck (decrease to 100 viral particles), necessitating that the initial viral generations of the next infection occur at low MOI. As in our study, Stern *et al*. conclude that viruses previously evolved at low MOI have a fitness advantage on a novel host. However, the mechanisms that mediate this advantage are likely be very different between the simulated poliovirus study and our Φ6 experiments. In particular, Stern *et al*. model host shift as a change in the fitness effects of a subset of nucleotide bases in the poliovirus genome, and they conclude that the advantage of low MOI evolved viruses is partly due to their ability to purge deleterious mutations rapidly. In our 24-hour assays, there is insufficient time for specific mutations to be fixed or purged in the population. The differences between L and H clones that we observed were more likely to be due to dissimilar phenotypes for host exploitation, such as faster adsorption (attachment) to cells to initiate infection, or faster lysis (burst) times to shorten the time between infection cycles.

We noted that the starting MOI in our current experiments (MOI = 10^−5^viral particles/cell) is much lower than low MOI employed in prior Φ6 studies (MOI = 0.002 viral particles/cell). While our extremely low initial MOI may have amplified host exploitation differences among viral clones, it is a reasonable value in the context of novel emerging diseases caused by RNA viruses. Natural MOIs within and across hosts and infection cycles are not widely established for most viruses, but transmission bottlenecks for some endemic and lab-studied diseases can be as low as a single or a few viral particles (Gutiérrez *et al*. 2012). Initial bottlenecks for emerging diseases are likely to be equally low. At the other end of the spectrum, MOI during cellular infection can range from < 0.01 particles/cell (in blood CD4+ T cells infected with HIV-1, calculated from Josefsson *et al*. 2011) to 13 particles/cell (for turnip infected with Cauliflower Mosaic Virus, Gutiérrez *et al*. 2010). Such large differences across hosts, viruses, and times of the infection cycle reveal that MOI – and the corresponding selective pressures – can vary widely across viral systems.

Within-host MOI may also affect which viral genotypes next experience transmission events. Some studies suggest that narrow transmission bottlenecks add stochasticity to viral evolutionary trajectories (Murcia *et al*. 2014). Our study in phage Φ6 suggests that intracellular competitiveness at high MOI trades off with replicative ability in other hosts. In turn, these results hint that viral genotypes evolved under high MOI may have a higher probability of transmission but may be less favored to replicate at the extremely low MOI conditions inherent to infecting the next host. On the other hand, a viral lineage may evolve such that certain genotypes (such as those advantaged under low MOI) are favored for transmission. For example, evidence from HIV studies suggests that certain genotypes may be selectively favored for their ability to be transmitted (Boeras *et al*. 2011).

MOI in natural viral populations is unlikely to be always high or always low. Rather, viral populations should be expected to experience stages of high MOI, followed by periods (at least occasionally) of low MOI. Future work should elucidate how these dynamic MOI cycles might affect long term adaptability of RNA virus populations, an idea that has been infrequently studied (e.g., Wilke *et al*. 2004). Here we document differences in the ability of a virus to infect novel hosts after evolution at two extremes of the MOI spectrum. Our findings suggest that MOI dynamics may play an underappreciated role in virus utilization of novel hosts.

## Acknowledgements

We thank C. B. Ogbunugafor, R. C. McBride, and J. W. Shapiro for their advice on experimental design and execution; R. Montville for providing supplemental data on phage burst assays; and Y. Salinas and members of the B. Kerr laboratory group for helpful comments and suggestions on the manuscript. We thank M. Hughes (Manager of the Statistical Consulting Center, Miami University, Oxford, Ohio USA) and J. Reuning-Scherer for R coding resources. This work was funded by the NSF BEACON Center for the Study of Evolution in Action (grant # RC062075YU) and an NSF GRFP grant (grant # DGE-0718124 and DGE-1256082) to SS. The authors declare no conflicts of interest.

## Data Archiving

Raw data and R scripts for data processing and analysis are archived at 10.6084/m9.figshare.4299266.

## Literature Cited

Baize S., D. Pannetier, L. Oestereich, T. Rieger, L. Koivogui, N. Magassouba, B. Soropogui, M. S. Sow, S. Keita, H. Clerck, G. Dominguez, M. Loua, A. Traoré, M. Kolié, E. R. Malano, E. Heleze, A. Bocquin, S. Mély, H. Raoul, V. Caro, D. Cadar, M. Gabriel, M. Pahlmann, D. Tappe, J. Schmidt-Chanasit, B. Impouma, A. K. Diallo, P. Formenty, M. Van Herp, S. Günther. 2014. Emergence of Zaire Ebola virus disease in Guinea. New Engl. J. Med. 371:1418–1425.

Barré-Sinoussi F., J. C. Chermann, F. Rey, M. T. Nugyre, S. Chamaret, J. Guest, C. Gauguet, C. Axlerblin, F. Vezinet-Brun, C. Rouzioux, W. Rozenbaum, L. Montagnier. 1983. Isolation of a T-lymphotrophic retrovirus from a patient at risk for acquired immune-deficiency syndrome (AIDS). Science 220:858–871.

Boeras D. I., P. T. Hraber, M. Hurlston, T. Evans-Strikfaden, T. Bhattacharya, E. E. Giorgi, J. Mulenga, E. Karita, B. T. Korber, S. Allen, C. E. Hart, C. A. Derdeyn, and E. Hunter. 2011. Role of donor genital tract HIV-1 diversity in the transmission bottleneck. Proc. Natl. Acad. Sci. USA 108:E1156–E1163.

Burmeister A. R., R. E. Lenski, J. R. Meyer. 2016. Host coevolution alters the adaptive landscape of a virus. Proc. R. Soc B283:20161528.

Clark B.P, O’Donald. 1964. Frequency dependent selection. Heredity 19:201–206.

Deardorff E. R., K. A. Fitzpatrick, G. V. S. Jerzak, P. Y. Shi, L. D. Kramer, and G. D. Ebel. 2011. West Nile Virus experimental evolution in vivo and the trade-off hypothesis. PLoS Pathog. 7: e1002335.

Dennehy J. J., S. Duffy, K. J. O’Keefe, S. V. Edwards, P. E. Turner. 2013. Frequent coinfection reduces RNA virus population genetic diversity. J. Hered 104:704–712.

Dixon M. G., Schafer I. J.Ebola viral disease outbreak – West Africa, 2014. 2014. Morbidity and Mortality Weekly 63:548–551.

Duffy S., P. E. Turner, C. L. Burch 2007. Pleiotropic costs of niche expansion in the RNA Bacteriophage ɸ6. Genetics 172:751–757.

Duffy M. R, T-H. Chen, W. T. Hancock, A. M. Powers, J. L. Kool, R. S., Lanciotti, M., Pretrick M., Marfel, S., Holzbauer, C., Dubray, L., Guillaumot, A., Griggs, M., Bel, A., Lambert, J., Laven, O., Kosoy, A., Panella, B. J. Biggerstaff, M. Fischer, and E. B. Hayes. 2009. Zika virus outbreak on Yap Island, Federated States of Micronesia. New Engl J. Med. 360:2536–2543.

Faye O., C. C. M. Freire, A. Iamarino, O. Faye, J. V. C. Oliveira, M. Diallo, P. M. A. Zanotto, A. A. Sall de 2014. Molecular evolution of Zika virus during its emergence in the 20th century. PLOS Neglect. Trop D 8:e2636.

Ford B. E., B. Sun, J. Carpino, E. S. Chapler, J. Ching, Y. Choi, K. Jhun, J. D. Kim, G. G. Lallos, R. Morgenstern, S. Singh, S. Theja, J. J. Dennehy 2014. Frequency and fitness consequences of bacteriophage ɸ6 host range mutations. PLoS ONE 9:e113078.

Ginsberg M., J. Hopkins, A. Maroufi, G. Dunne, D. R. Sunega, J. Giessick, P. McVay, K. Lopez, P. Kriner, K. Lopez, S. Munday, K. Harriman, B. Sun, G. Chavez, D. Hatch, R. Schechter, D. Vugia, J. Louie, N. Pascoe, S. Penfield, J. Zoretic, V. Fonseca, P. Blair, D. Faix, J. Tueller, T. Gomez, F. Averhoff, F. Alavrado-Ramy, S. Waterman, J. Neatherlin, L. Finelli, S. Jain, L. Brammer, J. Bresee, C. Bridges, S. Doshi, R. Donis, R. Garten, J. Katz, S. Klimov, D. Jernigan, S. Lindstrom, B., Shu, T., Uyeki, X. Xu, and N. Cox. 2009a. Swine influenza A (H1N1) infection in twochildren – Southern California, March-April 2009. Morbidity and Mortality Weekly Report 58:400–402.

Gutiérrez, S., M. Yvon, G. Thébaud, B. Monsion, Y. Michalakis, and S. Blanc 2010. Dynamics of the multiplicity of cellular infection in a plant virus. PLoS Pathog. 6:e1001113.

Gutiérrez S. Y. Michalakis S. Blanc. 2012. Virus population bottlenecks during within-host progression and host-to-host transmission. Curr. Opin Virol 2:546–555.

Harris F. J. 1978. On the use of windows for harmonic analysis with the discrete Fourier transform. Proc. IEEE 66:51–83.

Horiuchi K. 1983. Co-evolution of filamentous bacteriophage and its defective interfering particles. J. Mol. Biol. 169:389–407.

Huang, A. S. D. Baltimore 1970. Defective viral particles and viral disease processes. Nature 226: 325–327.

Huang A. S. 1973. Defective interfering viruses. Annu. Rev. Microbiol. 27:101–118.

Josefsson L.M. S. King, B. Makitalo, J. Brännström, W. Shao, F. Maldarelli, M. F., Kearney, W. S., Hu, J., Chen, H., Gaines, J. W. Mellors, J. Albert, J. M. Coffin S. E. Palmer 2011 Majority of CD4+ T cells from peripheral blood of HIV-1-infected individuals contain only one HIV DNA molecule. Proc. Natl. Acad. Sci. USA 108: 11199–11204.

Jordan H. T. M. C., Mosquera H. Nair, A. M. FranceMarch-April 2009. Morbidity and Mortality Weekly Dispatches. 2009. Outbreak of swine-origin influenza A (H1N1) virus infection — Mexico 58: 1–3.

Lalić J., J. M. Cuevas, S. F. Elena 2011. Effect of host species on the distribution of mutational fitness effects for an RNA virus. PLoS Genet 7:e1002378.

Lancaster K. Z, J. K. Pfeiffer 2012. Viral population dynamics and virulence thresholds. Curr. Opin. Microbiol 15:525–530.

Levy, J.A., A. D. Hoffman, S. M. Kramer, J. A. Landis, J. M. Shimabukuro, and L. S. Oshiro 1984. Isolation of lymphocytopathic retroviruses from San Francisco patients with AIDS. Science 225: 840–842.

Lewontin R. C. 1958. A general method for investigating the equilibrium of gene-frequency in a population. Genetics 43:419–434.

McBride R. C., C. B. Ogbunugafor, and P. E. Turner 2008. Robustness promotes evolvability of thermotolerance in an RNA virus. BMC Evol. Biol 8: 231.

Montville R., R. Froissart, S. K. Remonld, O. Tenaillon, and P. E. Turner 2005. Evolution of mutational robustness in an RNA virus. PLOS Biol. 3:1939–1945.

Murcia P. R., G. J. Baillie, J. Daly D. Elton C. Jervis, J. A. Mumford, R. Newton C. R. Parrish, K. Hoelzer, G. Dougan, J. Parkhill, N. Lennard, D. Ormond, S. Moule, A. Whitwham, J. W. McCauley, T. J. McKinley, E. C. Holmes, B. T. Grenfell, and J. L. Wood Intra-and interhost evolutionary dynamics of equine influenza virus. 2010. J. Virol. 84:6943–6954.

Smouse P. E. 1976. Implications of density-dependent population growth for frequency-and density-dependent selection. Am. Nat. 110:849–860.

Stern, A., S. Bianco, M. T. Yeh, C. Wright, K. Butcher, C. Tang, R. Nielsen, and R. Andino 2014. Costs and benefits of mutational robustness in RNA viruses. Cell Reports 8:1–11.

Turner, P. E., and L. Chao 1998. Sex and the evolution of intrahost competition in RNA virus ɸ6. Genetics 150:523–532.

Turner, P. E., and L. Chao 1999. Prisoner’s dilemma in an RNA virus. Nature 298:441–443.

Turner, P.E., and L. ChaoEscape from prisoner’s dilemma in RNA phage ɸ6. Am. 2003. Nat. 161:497–505.

Turner, P. E., and S. Duffy 2008. Phage evolutionary biologyBacteriophage Ecology: Population Growth, Evolution, and Impact of Bacterial Viruses. Cambridge University PressCambridge147–176.

Turner, P. E., N. M. Morales, B. W. Alto, and S. K. Remold 2010. Role of evolved host breadth in the initial emergence of an RNA virus. Evolution 64: 3273–3286.

Turner, P. E., J. A. Draghi, and R. Wilpiszeski 2012. High-throughput analysis of growth differences among phage strains. J. Microbiol. Methods 88: 117–121.

Vale, P. F., M. Choisy, R. Froissart, R. Sanjuán, and S. Gandon 2012. The distribution of mutational fitness effects of phage ɸX174 on different hosts. Evolution 66: 3495–3507.

Williams, E. S. C. P., N. Morales, B. R. Wasik, V. Brusic, S. Whelan, and P. E. Turner 2016. Repeatable population dynamics among vesicular stomatitis virus lineages evolved under high co-infection. Frontiers in Microbiology 7:370.

Wasik, B. R., A. R. Muñoz-Rojas, K. W. Okamoto,K. Miller-Jensen, and P. E. TurnerGeneralized selection to overcome innate immunity selects for host breadth in an RNA virus. Evolution 70:270–281.

Wasik, B. R., and P. E. TurnerOn the biological success of viruses. Ann. Rev. 2013. Microbiol. 67:519–541.

Wilke, C. O., D.D. Reissig, and I. S. Novella 2004 Replication at periodically changing multiplicity of infection promotes stable coexistence of competing viral populations. Evolution 58:900–905.

Woolhouse, M. E. J. and S. Gowtage-Sequeria 2005. Host range and emerging and reemerging pathogens. Emerg. Infect. Dis 11:1842–1847.

